# Electrical activity recorded from the spinal cord in freely moving rats using a subdural bioelectronic implant

**DOI:** 10.1101/2021.08.21.457239

**Authors:** Bruce Harland, Zaid Aqrawe, Maria Vomero, Christian Boehler, Brad Raos, Maria Asplund, Simon J O’Carroll, Darren Svirskis

**Author notes:** These authors contributed to the work equally (Bruce Harland, Zaid Aqrawe).

## Abstract

Bioelectronic devices have found use at the interface with neural tissue to investigate and treat nervous system disorders. Here, we present the development and characterization of a thin flexible bioelectronic implant inserted over the thoracic spinal cord in rats directly in contact with the spinal cord. There was no negative impact on hind-limb functionality nor any change in the volume or shape of the spinal cord. The bioelectronic implant was maintained in rats for a period of 3 months. We present the first subdural recordings of spinal cord activity in freely moving animals. Recordings contained multiple distinct voltage waveform shapes that were typically between 1 – 6 mV and lasted between 0.1 and 1 seconds. In the future, this implant will facilitate the identification of biomarkers in spinal cord injury and recovery, while enabling the delivery of localized treatments.

## Introduction

Bioelectronic devices have made significant contributions to the treatment of deafness, paralysis, epilepsy, and blindness through electrical recording and targeted stimulation of electrically active nerves (*1–4*). These devices are usually comprised of metallic electrodes which are patterned onto, and insulated by, flexible materials such as polyimide, parylene-C or poly- (dimethyl siloxane) (PDMS) (*5–7*). The area of application and intended use defines the final device geometry, electrode dimensions, and materials used. In this research, we investigate the use of a polyimide based bioelectronic implant to interface with the highly sensitive spinal cord environment, capable of monitoring spinal cord electrical activity and in the future facilitating electroceutical treatments directly to the injured spinal cord.

Polyimide and PDMS have both previously been described for use in contact with the spinal cord (*8, 9*). A PDMS spinal implant (e-dura) has been used to treat spinal cord injury (SCI) based on a different approach of neurostimulation to externally drive locomotor function in rats by bypassing the injury site (*9*). Targeted stimulation of the lumbosacral spinal cord during weight assisted treadmill rehabilitation led to improvements in locomotion during a 6-week period in rats with SCI. This treatment approach has been successfully translated to a clinical setting, resulting in recovery of voluntary control of leg muscles, which in some cases persists even after electrical stimulation was turned off (*10*). The e-dura, composed of stretchable PDMS, was inserted below the dura mater enabling electrical stimulation without the voltage drops associated with epidural placement, and allowing direct access to the spinal cord for drug administration (*10*). Compared to PDMS, a major advantage of polyimide is that the photo-lithographic methods for microfabrication are well-established. With metallizations patterned at micrometer precision using lift-off techniques, and with probe contours defined by dry-etching, devices can be made extremely compact, and still be durable and able to sustain long implantation periods (*11*). Polyimide had been used previously for epidural spinal implants (*8, 12*) but was found to be incompatible with subdural insertion, resulting in a profound foreign body response, deformation of the spinal cord, and impaired motor function compared to the PDMS implant (*9*). While the Young’s modulus of polyimide is far from the soft qualities of PDMS, the use of ultra-thin polyimide films has recently been demonstrated to achieve devices sufficiently compliant to conform to the contours of soft nervous tissue (*13*). With these recent developments in mind, we here re-explore the compatibility of an ultra-thin polyimide implant with subdural insertion along the thoracic spinal cord.

We show that our polyimide based bioelectronic implant is capable of recording spinal cord electrical activity in awake freely moving animals. In the future, this will allow electrical activity to be characterised in non-injured rats and SCI rats. This will increase the understanding of normal spinal cord electrical activity and changes that occur during injury and recovery. Meanwhile, the implant described is also capable of administering electroceutical treatments via the electrodes. Highly-promising laboratory data indicates electrical stimulation can regenerate injured neurons in cell-based studies (*14–16*), and this approach holds great promise for treatment of SCI. In the future, the bioelectronic implant will use elecrtrical recordings to ‘map’ the boundaries of an injury site and has tremendous potential for use in a clinical setting to detect the extent of injury and inform the location for delivery of personalised treatments.

To be an effective platform to probe spinal cord injury and deliver local electroceutical treatments, it is essential to show that i) the implant causes no impairment of motor function, ii) has minimal effect on spinal cord shape and foreign body response, iii) that it can be implanted for 3 months, sufficiently long to influence regeneration and then observe any resulting functional improvements after SCI (*17*), and iv) is capable of recording spinal cord electrical activity in an awake freely moving subject. Here, we describe the fabrication and characterisation of two designs of the bioelectronic implant, a planar version, and a bifurcated version to reduce pressure on the central spinal vein to alleviate potential behavioural or structural deficits post-implantation. We show that i) animals implanted for up to five weeks were healthy with no hind limb impairment, ii) implantation over one week had minimal effect on spinal cord structural integrity and foreign body response, iii) an external housing for the implants plug connector attached to back muscle via sutures and surgical mesh, allowed us to maintain the implant over a clinically relevant period for SCI treatment of 3 months, and iv) our implant can record electrical activity, which are, to our knowledge, the first spinal cord subdural electrical recordings from awake freely moving animals.

## Results

### 1. Electrochemical characterisation of the bioelectronic implant

Polyimide based bioelectronic implants were fabricated at a thickness of 8 μm based on two designs, planar and bifurcated (**Figure 1A**). Both designs had a total head width of 1.85 mm with 11 electrodes on each side, for a total of 22 electrodes, each with a diameter of 60 μm (electrodes numbered 1-11 on one side, and 17-27 on the other side (Figure 7 and Supplementary Figure 1)). The key difference between designs being the split down the middle of bifurcated devices, which was intended to reduce pressure on the central vein that runs along the dorsal midline of the spinal cord. Baseline electrochemical testing of the electrodes was undertaken in the form of cyclic voltammetry (CV) and electrochemical impedance spectroscopy (EIS). As first validation, and before implantation, each electrode was electrochemically characterised using cyclic voltammetry (CV) and electrochemical impedance spectroscopy (EIS) according to standard protocols (*18*). Representative CV scans of platinum microelectrodes embedded on either bifurcated or planar devices are displayed in **Figure 1B**. The scans show shapes that are typical of platinum microelectrodes, on both planar and bifurcated devices, with reversible reduction and oxidation peaks at −0.3 V and 0.4 V, respectively (*19*). Likewise, the EIS analysis in **Figure 1C** displays spectra typical of planar metallic microelectrodes, where the impedance throughout the frequency range of interest (1 Hz to 10,000 Hz) is dominated by the double-layer capacitance as visualised by a phase close to −90° and a slope on the |Z| vs frequency plot of −1 (*20, 21*). These baseline characterisation measures are important to assess the functionality of the entire system prior to implantation, from the implantable electrodes to the recording hardware.

**Figure 1:**
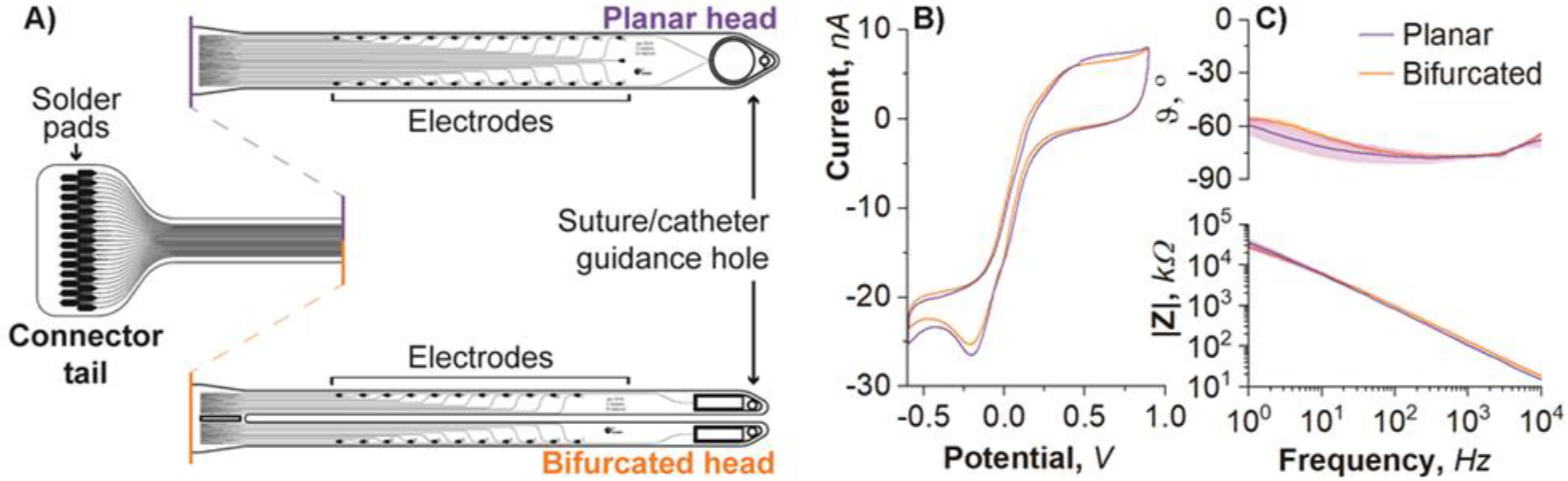
Two different configurations of the bioelectronic implant were designed and electrochemically characterised. (**A**) The bioelectronic implants consisted of a connector tail which comprised solder pads, each connected to tracts running down the body of the implant. Two ‘head’ designs were trialled and compared, being the planar head and bifurcated head. The platinum microelectrodes were characterised electrochemically in phosphate buffered saline through (**B**) cyclic voltammetry (representative result plotted) and (**C**) electrochemical impedance spectroscopy (n = 3 microelectrodes, data plotted as mean ± SD).

### 2. In vivo implantation and biocompatibility

The ultimate goal of our study is to develop a bioelectronic device capable of recording and stimulating the injured spinal cord over time periods relevant for spinal cord regeneration. For this to be possible, we first need to show that implantation of the polyimide based bioelectronic device does not negatively impact the structural integrity of the spinal cord, or the locomotor function. Furthermore, the implant must be well tolerated by the tissue over time, inducing minimal foreign body response. Furthermore, a direct comparison was made across two designs, planar and bifurcated (**Figure 1A**), to illicit whether the presence of a bifurcated head designed to reduce pressure on the central vein influences behavioural or structural differences post-implantation. To study this, spinal processes T12, T11, and T10 were removed to expose the dorsal surface of the thoracic spinal cord (**Figure 2A**) and planar or bifurcated implants were inserted subdurally (**Figure 2B**) in a cohort of rats for one week. Three different tests were carried out (i) behavioural assessment of rats post-implantation using the Basso Beattie Bresnahan (BBB) scale, (ii) structural assessment of the spinal cord directly below the position of the bioelectronic implant and (iii) assessment of neuroinflammatory responses through the quantification of astrocytes and microglia in spinal tissue directly below the position of the bioelectronic implant. In these experiments, the bioelectronic head was inserted along the spinal cord and the body of the implant positioned subcutaneously but severed from the printed circuit board and plug connector, so that no parts of the implant remained external to the animal. This allowed the impact of the spinal implant to be assessed without the influence of additional forces associated with an external connector.

**Figure 2:**
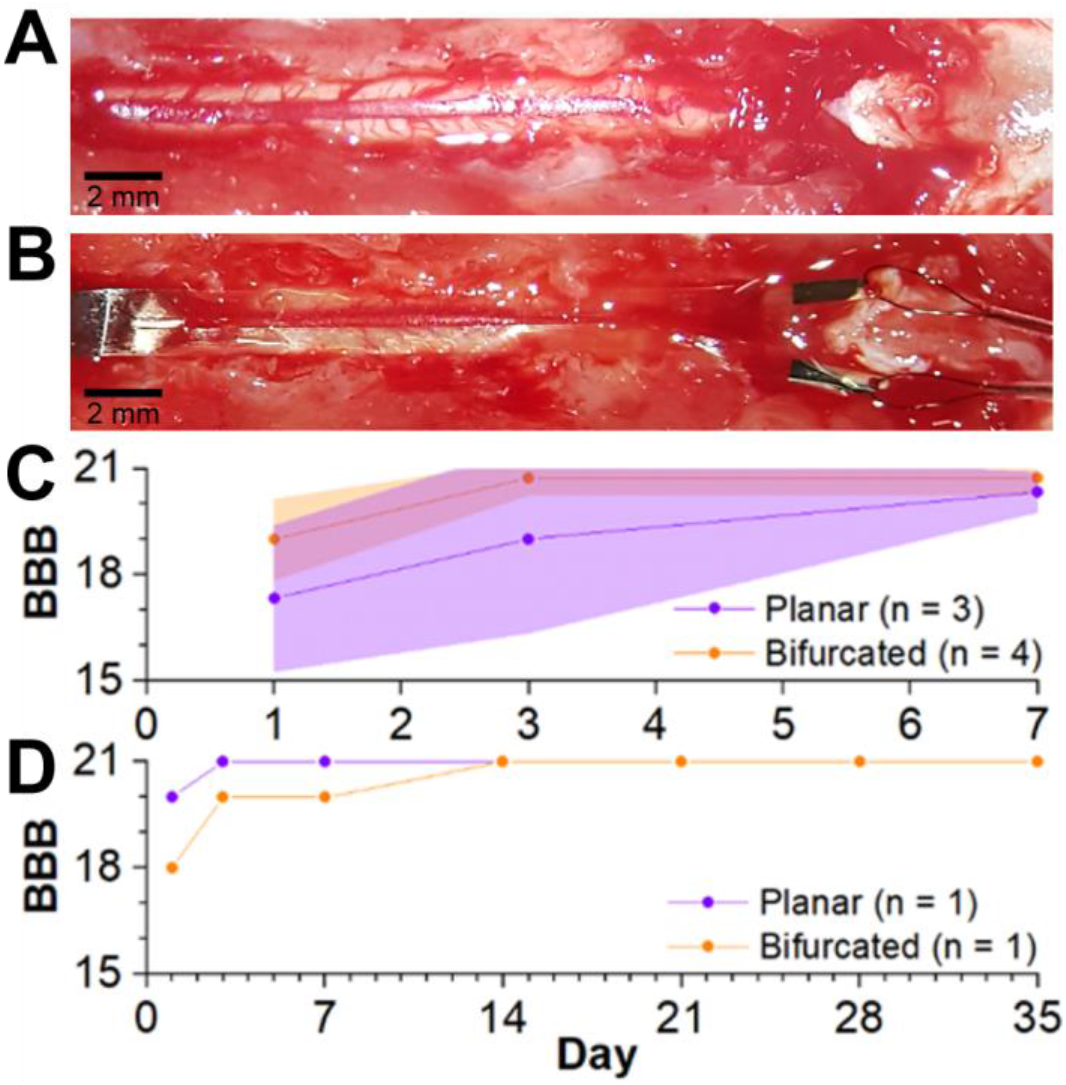
The bioelectronic implant had minimal impact on hindlimb function for up to five weeks after subdural implantation in rats. (**A**) Three spinal processes are removed (T12, T11, T10) and a section of spinal cord revealed. (**B**) A bifurcated implant is shown; each arm has been inserted into the subdural space from the left side of the image, and the tips of each arm exit the subdural space on the right side of the image. (**C**) Rats implanted with either version of the implant had only a transient loss of hind limb function and were generally unimpaired after a week (data reported as mean ± standard deviation). (**D**) Two of the animals (1 with bifurcated, and 1 with planar implant) were monitored for 5 weeks after implantation and showed no longer-term hindlimb impairment or other physiological issues.

#### 2.1. Insertion of the bioelectronic implant did not affect hind-limb function

Rats implanted with either planar or bifurcated devices displayed a mild transitory hindlimb deficit over the first week, with no statistical difference noticed between the two different implants, as measured through the BBB scale (**Figure 2C**). In all rats, there was a noticeable improvement in BBB-scores over time (Main Effect of Day; *F*(2,10) = 10.01, *P* < 0.01). Most recovery occurred by day 3 (Tukey’s post hoc: day 1 vs day 3, *P* < 0.05), and this remained until day 7 (day 3 vs day 7, *P* = 0.56). There was no main effect of Group (*P* = 0.19) or interaction (Group*Day; *P* = 0.43) which suggested that there was no difference between the bifurcated and planar implants in terms of hindlimb recovery. In both groups, animals were generally unimpaired at seven days post-implantation; all rats had scores of 21 meaning no impairment, or 20 meaning slight trunk instability but otherwise normal function (*22*). However, to ensure that a longer-term implantation did not result in decline in motor function beyond this weeklong recovery period, one animal with a bifurcated and one animal with a planar implant were retained for a 5-week period, and assessed with the BBB-scale at 14, 21, 28, and 35 days post-implantation. Both animals showed no impairment with consistent scores of 21/21 across this extended period (**Figure 2D**), in contrast to a previous report of moderate hindlimb deficits after insertion of a polyimide implant (*9*). After perfusion of the animals at Day 7, the implants were observed to still be in position along the spinal cord.

#### 2.2. The bioelectronic implant did not affect spinal cord volume or shape

Although the bioelectronic implant had no long-term impact on motor function, we assessed the effect of the bifurcated and planar implants on spinal cord segments directly underneath the implants (T13, L1, and L2) at 7-days post-implantation (**Figure 3A**). It is important to note that spinal segment labels do not correspond to the names of the spinal processes positioned directly above them (*23*). In a previous study, insertion of a 25 μm thick polyimide implant (*cf*. 8 μm thick implants used in the current study) resulted in a severe deformation in spinal cord volume and shape below the implant (*9*). We collected coronal spinal sections, stained with haematoxylin and eosin to aid with visualisation of the tissue (**Figure 3B**) from rats implanted with bifurcated (n = 3) and planar (n = 2) implants for a week and compared them with sections obtained from control rats (n = 3), which received the laminectomy only. For each section, the perimeter was traced to obtain the surface area (cm^2^) and shape (roundness).

**Figure 3:**
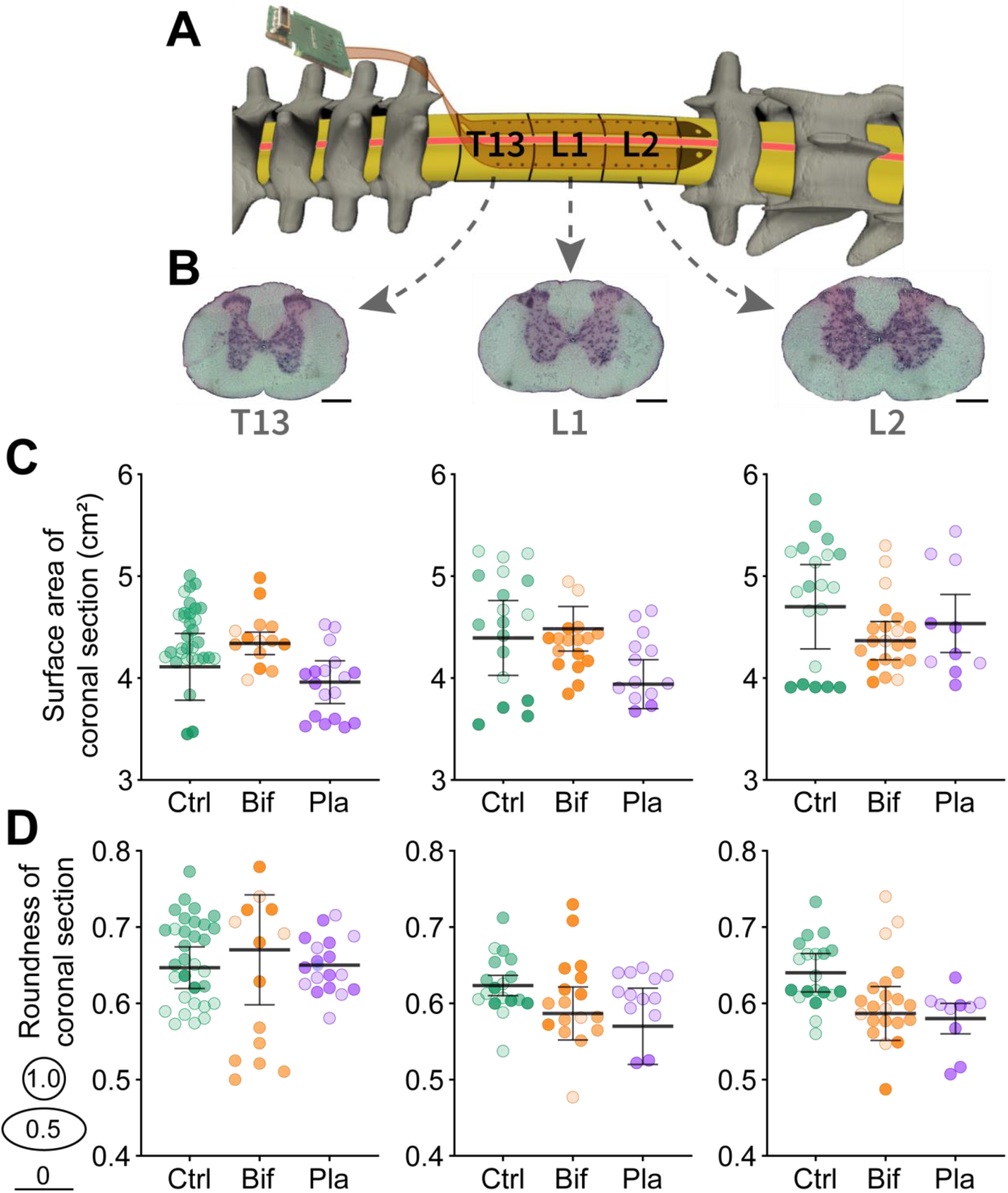
The bioelectronic implant did not significantly impact spinal cord volume or shape. (**A**) Spinal cord segments directly underneath the bioelectronic implant were examined. Note that spinal segment labels do not correspond to the names of the spinal processes above them (*23*). (**B**) Examples of T13, L1, and L2 coronal sections of spinal cord stained with a haematoxylin and eosin stain to aid with visualisation of the tissue are shown (scalebars = 0.5 mm). The perimeter of each section was traced to assess the surface area and shape. (**C**) The surface area of coronal sections is shown, taken from rats without implants (Ctrl), planar implants (Pla) and bifurcated implants (Bif) at the T13 (left), L1 (centre) and L2 (right) segments of the spinal cord. (**D**) The shape of sections was also assessed via a roundness score between 1 (circular) and 0 (flat line) at T13, L1 and L2 sections (from left to right) of the spinal cord. Coronal sections are shaded differently for each individual animal on the dot plots, error bars show standard error between animals. Comparisons using one-way Anova’s revealed no group differences.

The average surface area of coronal sections was not significantly different between groups when compared with ANOVA for any of the spinal segments (T13: *F*(2,5) = 0.58, *P* = 0.59, L1: *F*(2,5) = 0.8, *P* = 0.5, L2: *F*(2,5) = 0.3, *P* = 0.75). Coronal segments from the different groups were observed to be similar to each other with no noticeable reduction in size associated with the presence of the implant. However, there was a non-significant trend for planar implant sections to be smaller at T13 and L1 (**Figure 3C**).

The shape of spinal cord sections was assessed using a roundness score, which ranged from 1, indicating circular, to 0, indicating a flat line. ANOVAs indicated no significant difference between the groups in the shape of the spinal segments (T13: *F*(2,5) = 0.06, *P* = 0.94, L1: *F*(2,5) = 0.7, *P* = 0.54, L2: *F*(2,5) = 1.25, *P* = 0.26). In general, the shape of spinal coronal sections were similar between groups and there were no malformations as were previously reported (*9*). However, there was a non-significant tendency for sections to be slightly less round in the presence of either implant compared with the controls at L1 and L2 (**Figure 3D**).

We investigated whether deformation of the cord occurred over a more extended period in two of our rats that were implanted for five weeks across two spinal cord segments (L1 and L2). After five weeks, spinal segments below the implants had an average roundness of 0.68 ± 0.07 at L1 and 0.62 ± 0.03 at L2 compared with the one-week control animals, which had roundness scores of 0.62 ± 0.04 at L1 and 0.64 ± 0.04 at L2. The 5-week implanted rats had rounder L1 spinal segments than controls and shape did not differ at L2. This difference in shape may be related to the increased size and weight of these 5-week implanted rats compared to the 1-week controls (~15% increase in average weight) and we did not compare surface area between these groups for this reason.

#### 2.3. Foreign body response related to the presence of the implant was limited

Implantation of devices on the surface of the brain or spinal cord typically results in a foreign-body neuroinflammatory response of macrophages (microglia) and astrocytes below the position of the implant (*24–26*). Therefore, coronal sections from two animals per group were immunohistochemically stained with standard markers for astrocytes (GFAP) and microglia (Iba1). In order to focus on changes in these markers directly associated with the presence of the implants on the dorsal surface of the spinal cord, only the region of spinal tissue directly below the implant was isolated (**Figure 4**). Comparison with the control animals, which received the laminectomy surgery but no implant, provided a measure of foreign-body response associated with the week-long presence of the bifurcated and planar implants.

**Figure 4:**
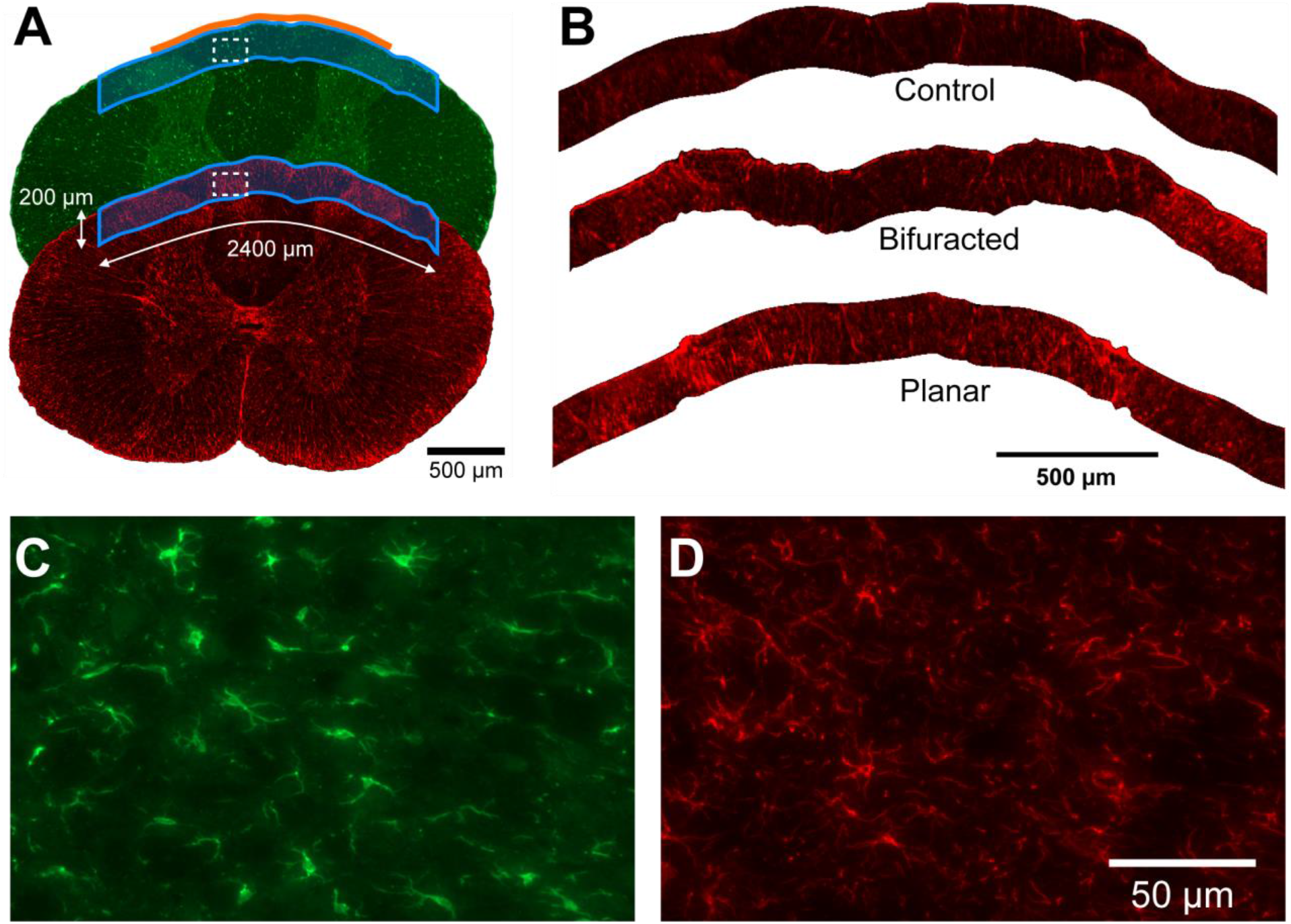
The neuroinflammatory response of astrocytes and microglia was examined in spinal tissue directly below the position of the bioelectronic implant. (**A**) Coronal spinal cord sections were stained with standard cellular markers for astrocytes (GFAP; red) and microglia (Iba1; green). The approximate position of the bioelectronic implant is shown by the orange line at the top of the image (not to scale). For each section, only the region of interest (ROI) of spinal cord tissue directly below the implant was isolated and analysed (blue boxes). Sections shown are from a rat implanted with planar implant. (**B**) Representative examples of ROI’s from each group stained with GFAP to label astrocyte-response (example ROI’s are from spinal segment T13). Examples of (**C**) microglia stained with Iba1, and (**D**) astrocytes stained with GFAP are shown at 40x magnification, these images were taken from the white dotted box indicated in panel A.

The fluorescence intensity of labelled astrocytes was significantly different between the groups at spinal segment L2 but not at T13 and L1 (ANOVA’s: for L2, *F*(2,3) = 16.73, *P* < 0.05; for T13, *F*(2,3) = 2.46, *P* = 0.11; for L1, *F*(2,3) = 3.63, *P* = 0.16; **Figure 5A**). At L2, the group difference was due to an increased astrocyte response in planar implanted rats compared to control rats (Tukey’s post hoc: *P* < 0.05). Florescent intensity of microglia was not significantly different between the groups for any of the spinal segments (ANOVA’s: for T13: *F*(2,3) = 1.94, *P* = 0.29), for L1 (*F*(2,3) = 0.75, *P* = 0.54; for L2: *F*(2,3) = 1.33, *P* = 0.39; **Figure 5B**).

**Figure 5:**
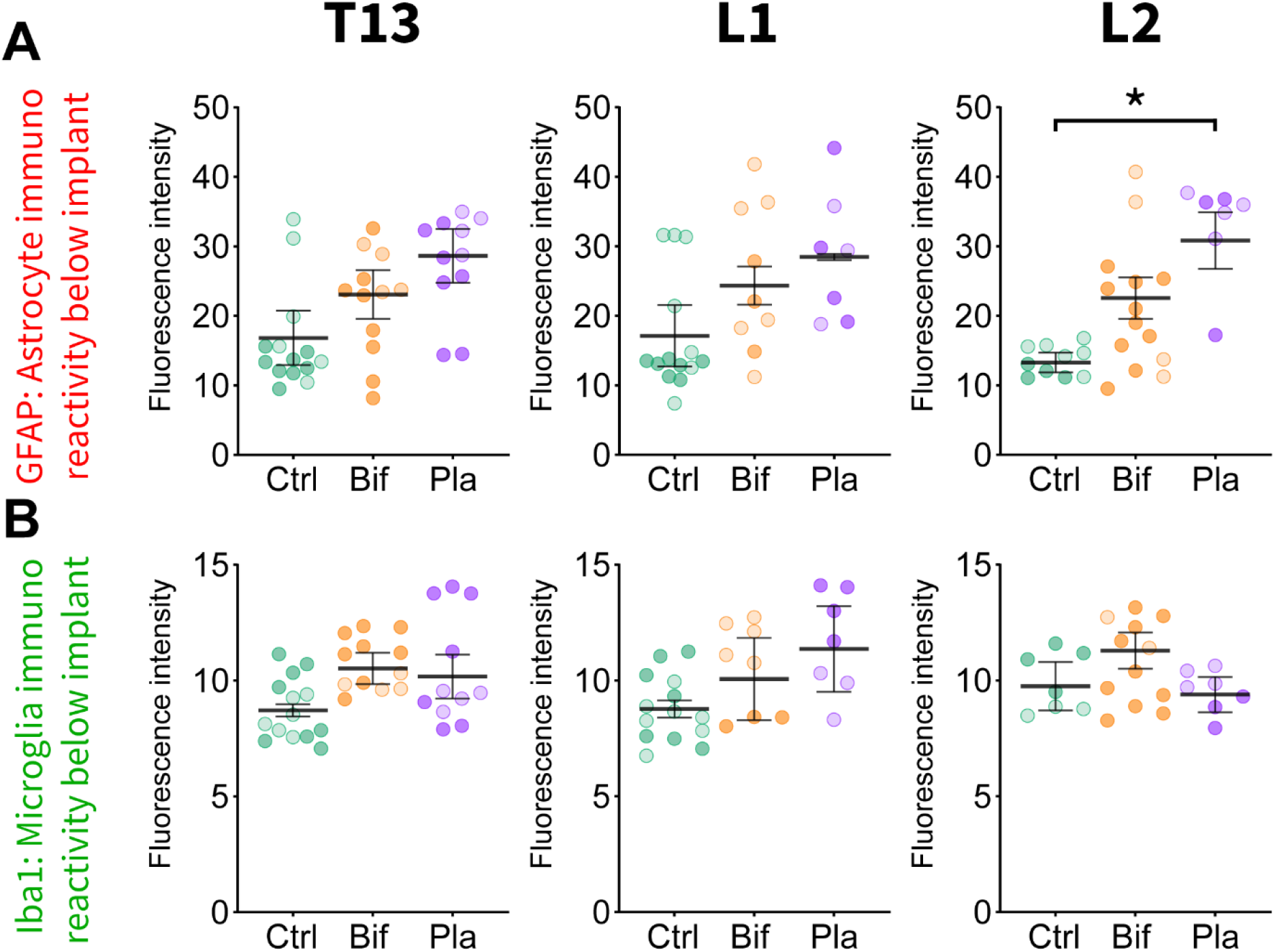
The planar implant had a localised foreign body response of astrocytes compared to controls. (**A**) The average fluorescence intensity of astrocytes stained with GFAP is shown for ROI’s across the three thoracic segments for the controls (Ctrl), bifurcated (Bif), and planar (Pla) implants. Rats implanted with the planar implant had greater activation of astrocytes compared to the control group in the L2 spinal segment only. (**B**) Rats implanted with either implant did not differ from controls in terms of Iba1 labelled microglia expression. ROIs are shaded differently for each individual animal on the dot plots, error bars show standard error between animals. All comparisons were one-way ANOVA’s, * = *P* < 0.05.

### 3. Electrophysiological spinal cord recordings

#### 3.1. The implant and external plug housing remained in position for up to 3 months

Four rats had bifurcated bioelectronic implants inserted along the thoracic spinal cord, and backpack assemblies (**Figure 6A**) were attached to the back muscles via sutures and surgical mesh. Animals were plugged in two or three times per week for 3 – 5-minute recording sessions, during which they ambulated and sat upon on a raised platform (**Figure 6B**). Backpacks remained viable and firmly attached to the back muscle and skin of the animals over a 3-month period (86, 89, 98, and 97 days). Two of the animals were tested in the open field using the BBB-scale, and both scored 21/21 at 1, 3, and 7 days after surgery, and every week after that, up to 12 weeks post-implantation. To examine implant positioning, one of the animals was perfused at 89 days post-implantation. The backpack and outer muscle was carefully removed, and the bioelectronic body of the implant was severed, leaving the implant head in position along the spinal cord. A cylinder containing the intact spinal cord and including ~1 cm of tissue at every side was dissected and processed for micro-CT scanning. When the scanned images were reconstructed, a dorsal ‘bird’s eye’ view showed both arms of the implant still positioned appropriately along the spinal cord inside the laminectomy (**Figure 6C**), which confirmed that the implant had not been displaced over the 3-month period.

**Figure 6:**
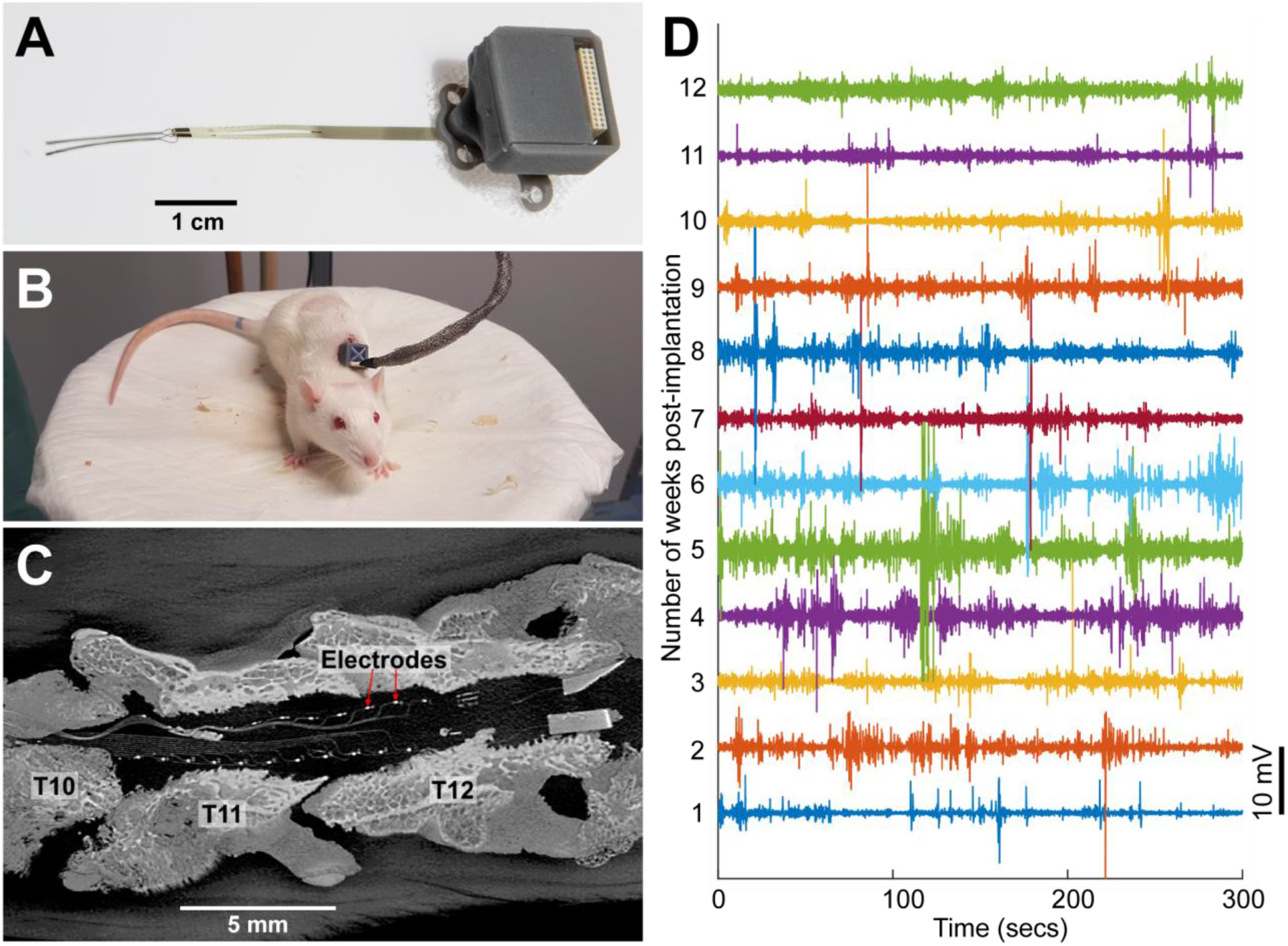
The bioelectronic head remained in position along the spinal cord after 3 months in rats implanted with a backpack housing for the external plug. (**A**) A resin printed backpack housed the printed circuit board (PCB) and plug. A circular piece of surgical mesh was attached to the ventral surface of the backpack, which was positioned subcutaneously forming a strong attachment to the tissue. (**B**) The backpacks allowed rats to be plugged in via a recording cable while sitting and ambulating upon a raised platform. (**C**) After 3 months, a cylinder of tissue was excised, containing the intact spinal cord around the bioelectronic head of the implant, and micro-CT scanned to ascertain the position of the implant. An overhead view of the bioelectronic head shows that the electrodes still oriented correctly along the surface of the spinal cord (red arrows point to two electrodes). The scour marks in the spinal bone where the T10, T11, and T12 spinal processes were removed are visible, showing the implant is still in the expected rostral-cranial position. (**D**) Spinal cord electrical activity was recorded while the rat was on the raised platform, recordings from the same electrode in one of the rats is shown over 12 weeks.

#### 3.2 Spinal cord electrical activity was recorded in freely moving rats

Recordings of spinal cord electrical activity were taken 2 – 3 times a week throughout the full study period of three months. **Figure 6D** shows an example of weekly recordings from a single electrode midway down the left bifurcated arm of the implant; which was positioned on the right hemisphere of the spinal cord above spinal segment L1. The electrical recordings remained viable over the 12-week time-course, and no degradation of the signal was noticeable. **Figure 7** shows electrical activity from every electrode during a single recording, which was taken 7 days after implantation in one of the rats, the same time point used for the histology in the earlier experiments. The electrical activity is shown for the right and left arms of the bioelectronic implant. Within this recording session, the amount of activity varied across all of the electrodes, and a higher general level of activity is noticeable on one of the implant arms (right side of **Figure 7**). Furthermore, voltage spikes greatly ranged in their magnitude, which may indicate distance from the source of the electrical signal.

**Figure 7:**
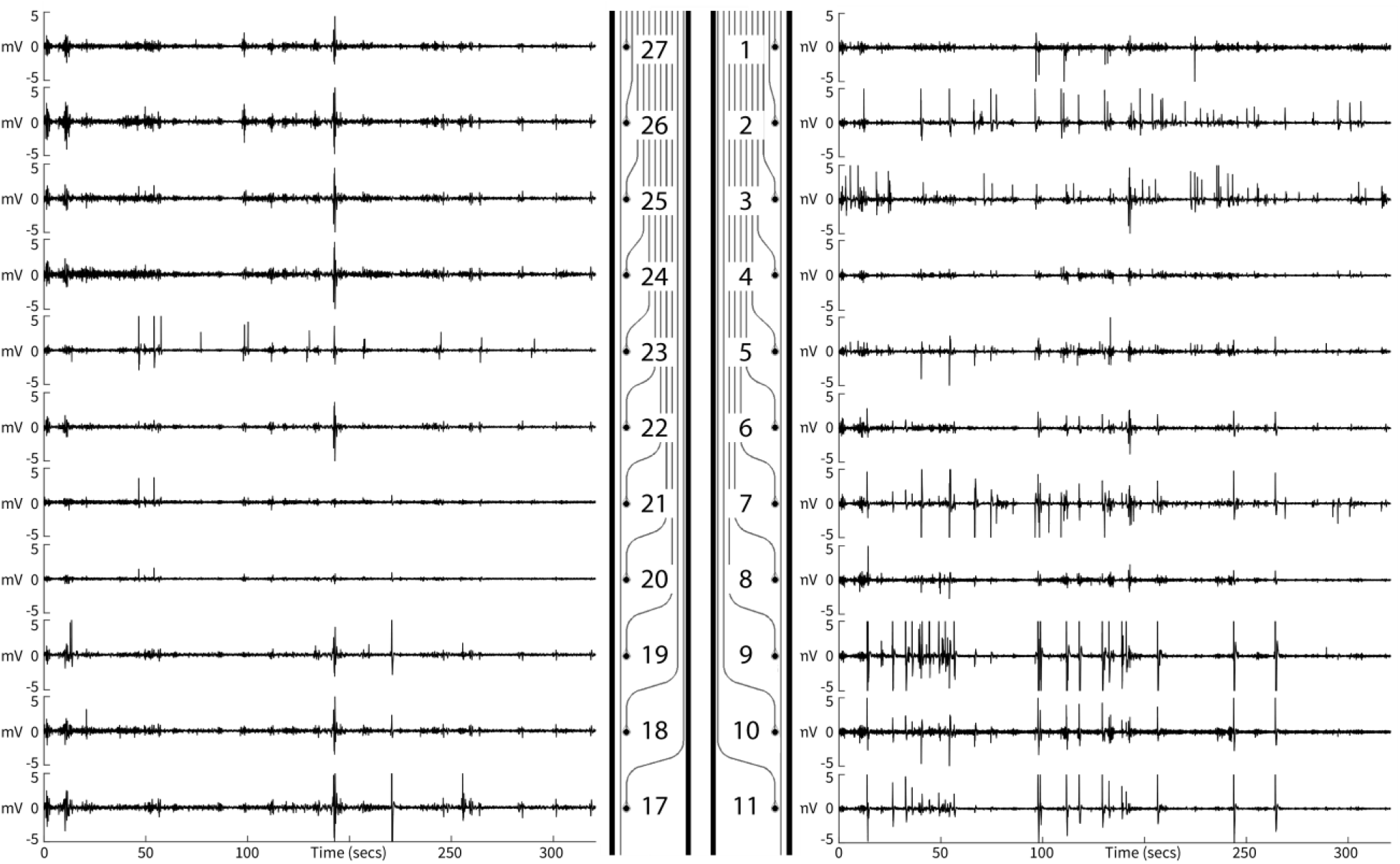
Electrical activity patterns varied across electrodes during recording sessions. Concurrent electrode activity is shown for the right and left arms of the bioelectronic implant while the rat ambulated on a raised platform for five minutes. The relative activity varies across electrodes, and voltage spikes vary greatly in magnitude which may indicate the distance from electrode to the electrical source in the tissue. Numbers on the implant diagram correspond to the electrode ID.

A 50 Hz notch filter was applied to the raw data to remove noise associated with the mains power. A median subtraction was then performed to remove noise and artefacts associated with movement of the animal and recording cable. **Supplemental Figure 1** demonstrates the effect of the median subtraction on the voltage recorded from four electrodes. The median subtraction removed changes in voltage that were common to every electrode and emphasized voltage changes that were unique to a single or small number of electrodes. The recordings were made as the rats moved around a raised platform and consisted of a range of activity including walking, sitting, rearing, and grooming. **Figure 8A** shows an enlarged image of a single electrode’s activity (electrode 10 from **Figure 7** and **Supplementary Figure 1**) recorded at 7 days post-implantation. The electrical activity varies greatly in magnitude with a few higher amplitude spikes evident in the range of 1 – 6 mV, whereas the majority of signals have an amplitude below 2 mV. These differences in spike magnitude may be related to varying distance between source and electrode (*27*). Three characteristic voltage waveforms were observed throughout the recording (**Figure 8B**). The first waveform (**Figure 8, B1**) consists of a sharp negative change in potential away from the baseline potential, a slower return that undershoots the baseline, followed by a slow return to the baseline potential. **Figure 8D** shows that magnitude of the initial voltage spike ranged from 1 – 6 mV and the total length of the waveform ranged from 0.5 to 1 seconds. Similarly, the second waveform (**Figure 8, B2**) has the same shape, but with the opposite polarity, consisting of an initial sharp positive deviation from the baseline potential. Both characteristic waveforms 1 and 2 were observed with a range of magnitudes and have been grouped in **Figure 8B** according to their absolute magnitude. Notably, several of waveforms 1 and 2 showed two successive initial voltage spikes of the same polarity. The differences in polarity between waveforms 1 and 2 may reflect the angle or direction of propagation of the signals, whether they arise from dorsal root ganglion afferents or comprise signals passing through from more rostral segments of spinal cord. The parallel rows of electrodes on the bifurcated arms would be positioned along the surface of the spinal cord at the border of the inside arch of the dorsal horn and the dorsal columns, a region containing both neurons and bundles of axons involved in relaying somatosensory signals from the hind-feet and hind-region of the body to the brain (*28*). The third characteristic waveform (**Figure 8, B3**) consists of smaller magnitude negative voltage spikes that reach between approximately 1 and 2 mV below the baseline but oscillate in rapid succession. **Figure 8E** shows that these repetitive voltage spikes generally occurred at a frequency of 7 – 10 Hz. However, all the described waveforms occur at a lower frequency than brain signals as has been reported in previous studies, which have recorded spinal cord activity in anesthetised rats (*9*), and *in vitro* spinal cord slices in rats and mice (*29, 30*).

**Figure 8:**
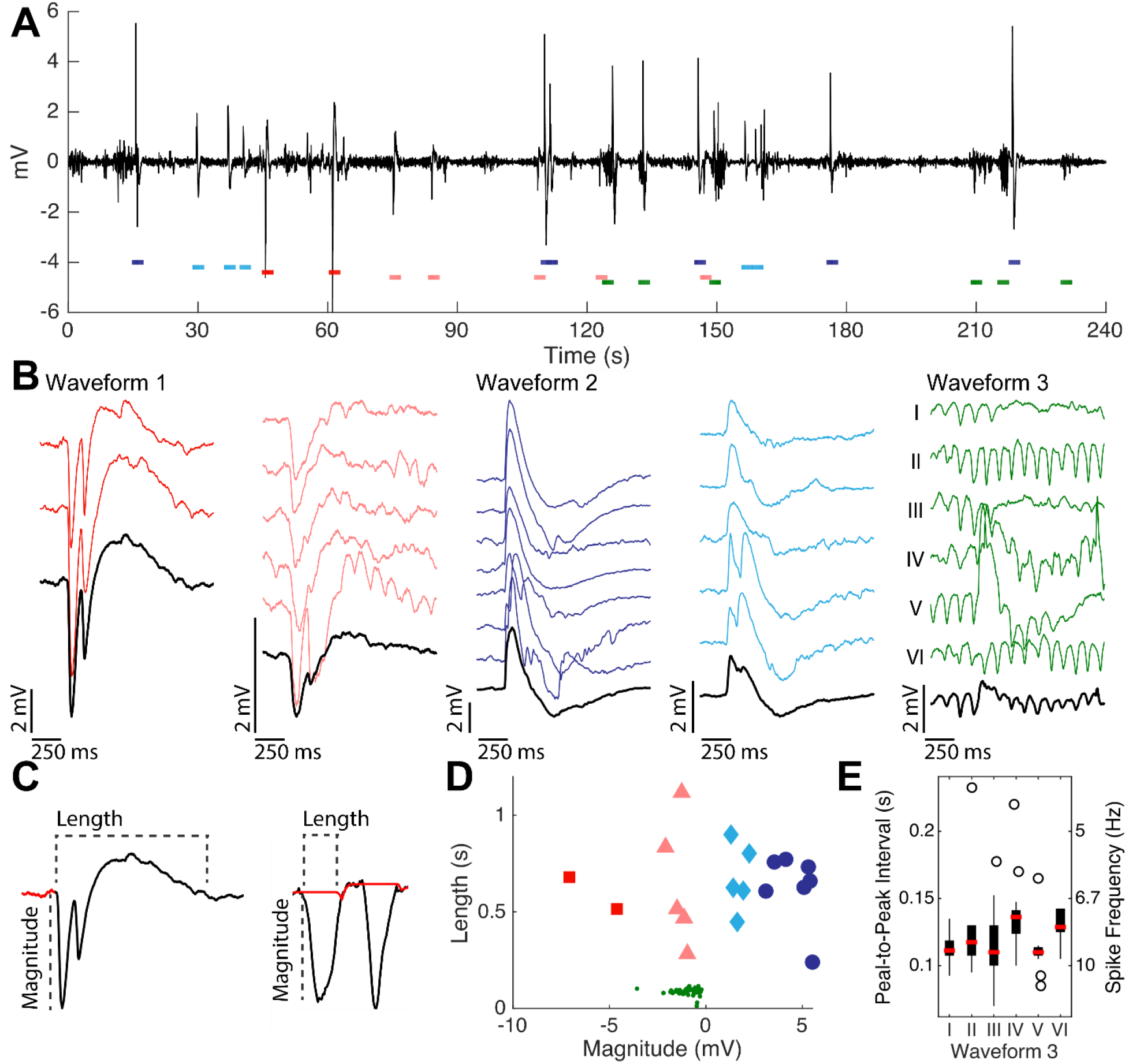
Spinal cord electrical activity recorded from a single electrode. (**A**) A four minute recording from a single electrode shows electrical activity across a range of amplitudes. Colored bars indicate locations and group of the voltage waveforms in B. (**B**) Three characteristic voltage waveform groups recorded from the spinal cord in unrestrained rats while ambulating and sitting upon a raised platform. Waveform groups 1 and 2 have been further split by absolute magnitude (0 – 2.5 mV and 2.5 mV and greater, respectively). Black waveforms represent the mean waveform within each subgroup. (**C**) Definition of waveform baseline (red), magnitude and length for single spike (B1 and B2) and oscillatory (B3) waveforms. See methods for details. (**D**) Point cloud of voltage waveform length and magnitude. Colors correspond to the waveforms in B. (**E**) For waveform group 3, the spike frequency is shown for the six individual waveforms.

## Discussion

This work details the fabrication and characterization of an innovative subdural bioelectronic implant designed to be positioned above a spinal cord injury to monitor changes in electrical activity and capable of acting as a delivery platform for electroceutical and chemical treatments. Insertion of the internal parts of the implant in rats had no effect on physiological function or spinal cord volume and shape, at least over the 5 weeks studied here. Foreign body response to the presence of the implant was only detectable when the planar design of the implant was used, and only localised to one segment of the spinal cord. Next, we designed an external housing for the implant capable of being surgically attached to the upper back muscle for a period of up to 3 months, after which we were able to show that the bifurcated implant remained in position along the spinal cord. Lastly, we demonstrate electrical recordings captured from the device in awake freely moving rats.

The bioelectronic implant is constructed of a thin flexible polyimide, which is a highly durable material that shows excellent compatibility with nervous tissue, which previously have been demonstrated for intracortical and epicortical microdevices (*6, 11, 13, 31–33*). However, a previous study showed that a polyimide implant inserted along the lumbosacral spinal cord in rats resulted in significant hindlimb motor deficits six weeks after surgery (*9*). Minev et al. found that polyimide implanted rats made significantly more errors on a horizontal ladder task than sham rats, and rats implanted with a PDMS implant. Furthermore, a comprehensive analysis of hindlimb gait revealed that polyimide implanted rats had significantly impaired foot control in terms of step height, orientation, and hip joint amplitude, as well as reduced leg movement, and lateral foot placement compared to sham and PDMS implanted rats. This suggests that these rats would have scored around 12-15 on the BBB-scale. In contrast, animals in the current study only showed some mild transient impairments 1 – 3 days after surgery, with no impairment after one week and the following weeks up to five weeks post-implantation. Minev et al. did not assess hindlimb function before the six-week mark, but it can be assumed that function was the same or worse at these time-points. The different outcomes in these studies are most likely explained by differences in the design of the polyimide implants used. The implant described by Minev et al. was just over three times thicker (25 vs 8 μm) and almost twice as wide (3.2 vs 1.85 mm) as the bioelectronic implants described here. It should be noted that in previous experiments we implanted wider and thicker versions of our bioelectronic implant in rats (2.15 mm width, 12 μm thick), which resulted in hindlimb deficits in some animals (data not included). It is also important to point out that in Minev et al., implants were positioned further back (spinal segments L2 to S1) and covered twice the length of spinal cord than in the current study (3 vs 1.5 cm). However, it is evident that the width, thickness and design of the implant play a key role in its tendency to cause damage to the spinal cord environment and result in physiological impairment whilst implanted. Thus, the difference in mechanical properties between the elastomer PDMS and polyimide are here compensated by the latter allowing for an ultra-thin design, which ensures structural biocompatibility of the resulting implant. This is well in line with previous work concluding that an 8 μm polyimide layer was sufficiently thin to allow an epicortical grid to conform to the curvature of the rat brain at no additional mechanical pressure (*13*). Our data suggests that 8 μm must be very close to the ‘ideal’ device thickness for achieving full conformability for the spinal cord for these animals (2 month old female Sprague-Dawley rats). Considering the added difficulties associated with PDMS implant manufacturing, such as higher impedance in connection lines and challenges with stretchable electrode materials, it is highly encouraging that similar structural biocompatibility appears to be possible using a non-stretchable polymer such as polyimide and conventional thin-film metallizations (*34*).

Impairments in locomotor function would be expected to be accompanied with changes to the volume and shape of the spinal cord in the implanted area (Minev et al. 2015). We found no differences in spinal cord volume or shape in rats implanted for one week, and no further changes in animals implanted for five weeks. In contrast, Minev et al. reported a severe deformation of the spinal cord in rats implanted with their thicker and wider planar shaped polyimide implant for six weeks, which was accompanied with hindlimb impairments (*9*). Although surface area was not quantitatively assessed, their representative 3D reconstruction showed that the spinal cord was significantly depressed along the entire region below their polyimide implant. This is likely due to the much greater thickness and width of their implant, which could lead to deformation of the cord over time in the limited subdural space (*13*).

We did not detect an increase in the presence of microglia in the region of spinal cord directly below the bioelectronics implant, although there was an increase in astrocytes in the L2 region for the planar design of the implant only. This difference may be a result of the planar style implant straddling the posterior spinal cord vein which is positioned above the surface of the spinal cord in the subdural space, whereas the bifurcated arms are positioned either side. Therefore, the planar device may exert more pressure upon this central vein (*35*) and would not conform as well to the spinal cord surface on either side. It is likely that the bifurcated implant design avoids these issues in the same way that highly fenestrated polyimide-based epicortical devices have been shown to conform better to surface of the brain (*13*). For this reason we used the bifurcated design exclusively for the longer-term electrophysiology experiments. Foreign body response due to the presence of the bioelectronic implant may be more pronounced over a longer time-span of implantation (*25*), which will be investigated in future work.

The backpack housing, positioned just below the shoulder blades on the upper back, is a modified versions of the lower back housings developed by Wurth et al. for sciatic nerve implants (*36*). The backpack has several advantages over skull mounting, which is used in brain neural recording devices (*37, 38*), and has also been used for spinal cord implants (*9, 12*). The use of the backpack allows for a less invasive surgery, as the device is positioned just in front and above the laminectomy in the same incision site used for insertion of the bioelectronic implant. This can be useful when combining the implant with a mild or moderate spinal contusion injury model, in which many changes in physiological function due to natural recovery occur in the first week after injury, which coincides with surgical recovery. Skull-mounted devices use micro screws embedded in acrylic cement to make a strong attachment, and the dorsal skull plates are often removed with the device. Our bioelectronic implant is designed to deliver electrical treatments to an injury site to regenerate damaged neural tissues, after which the implant could be removed. Unlike a skull-mounted device, the backpack can be surgically removed once recovery is achieved. However, the viability of the backpack anchoring method will need to be further tested in spinal cord injured rats and over the planned experiment time (3 months) with the inclusion of frequent recording and stimulation sessions, which will put additional strain on the device. We envisage that 3 months is sufficiently long for treatments to influence regeneration and any resulting functional improvements to be observed in a rat model of SCI (*17*).

The bioelectronic implant has tremendous potential for future use in testing treatments for SCI, both electroceuticals (electric field therapies) and as a drug delivery platform capable of applying regenerative treatments directly to the injured spinal cord for sustained periods. SCI is a devastating condition that can result in permanent neurological impairment, and there are currently no regenerative therapies available (*39*). However, promising approaches to achieve axonal regeneration have been identified *in vitro*, such as the application of electrical fields (*14*). Weak electric stimulation (up to 80 mV/mm) has been shown to enhance axonal regeneration as well as guide orientation *in vitro* (*16*). However, a major challenge is to apply treatments directly to a SCI and over a continuous treatment period. Although not explicitly addressed in this work, the subdural polyimide bioelectronic implant we describe here is suitable for providing spatiotemporally precise electric field generation delivered via its electrodes. As the fabrication follows standard protocols for thin-film bioelectronics, functional electrode materials such as nanostructured platinum, iridium oxide or PEDOT/PSS could be added in a few additional steps according to previously described protocols (*19*).

Here we have shown spinal cord electrical recordings in freely moving rats for the first time, over a period of twelve weeks. In future work we plan to characterize this activity over a 3-month period and compare uninjured control rats with SCI rats, and SCI rats treated with daily electrical stimulation across the injury site. Subtle changes in electrical activity after SCI may provide early indications that treatments are working. Furthermore, using electrical recordings to map the boundaries of an injury site could also have additional value in a clinical setting to detect the extent of an injury and inform personalised electroceutical treatments.

We have demonstrated that our newly developed bioelectronic implant, which can be safely implanted along the spinal cord for a clinically relevant period, has no deleterious effect on motor function and only minimal effect on spinal cord shape and foreign body response. Therefore, the device has great potential to monitor electrical signaling in the spinal cord after an injury, and in the future to deliver localised treatments for spinal cord injury.

## Methods

### Bioelectronic Implant Fabrication

Two different polyimide device designs were fabricated comprising a (i) planar 1.85 mm width and (ii) bifurcated 1.85 mm width polyimide device (**Figure 1**). These widths were chosen as the diameter of the spinal cord was approximately 3 mm. The bioelectronic devices were fabricated using standard microlithography techniques in the cleanroom facility at University of Freiburg (RSC, ISO 5, according to ISO 14644-1). First, U-Varnish-S polyimide (PI, UBE Industries, Ltd, Japan) was spun on silicon wafers (at 3000, 4500 and 9000 rpm for achieving an after-annealing thickness of 6, 4 and 2 μm, respectively). Then, tracks/interconnection lines, connection pads and active sites were patterned using the high resolution image reversal photoresist AZ 5214 E (MicroChemicals GmbH, Germany). O2 plasma (80W, Plasma System 300-E, PVA TePla, Germany) was used to activate the surface and as an adhesion promoter for the metal, which was subsequently evaporated onto each wafer (100 nm of Pt, Univex 500 Electron-Beam Evaporator, Leybold GmbH, Germany). Lift-off to remove excess metal was done in acetone. O2 plasma was again used to activate the surface of the wafers before a second PI layer was spun onto all of them. The positive photoresist AZ 9260 (MicroChemicals GmbH, Germany) was used as a masking layer for the opening of the contact pads and electrode sites and for setting the outlines of the devices. PI was finally etched using O2 plasma in a reactive ion etching (RIE) machine (STS Multiplex ICP, SPTS Technologies, United Kingdom) with a multi-step protocol (200 W first and 100 W after, time varying with PI thickness). The photoresist was stripped off in an acetone bath and the devices were mechanically peeled off from the wafers using flat tweezers.

### In vitro Electrochemical Characterisation

Prior to implantation, all electrodes on respective devices were characterised electrochemically through electrochemical impedance spectroscopy (EIS) and cyclic voltammetry (CV) measurements in a phosphate buffered saline (PBS, 0.01 M) electrolyte. An electrochemical workstation (Biologic) with a three-electrode set-up was utilised and comprised a silver/silver chloride (Ag/AgCl) reference electrode, platinum counter electrode and the respective platinum electrode to be tested on the bioelectronic device as the working electrode. For CV measurements, potential limits were set at 0.9 V and −0.6 V vs Ag/AgCl and each electrode was scanned three times at a scan rate of 100 mV s^−1^. EIS measurements were carried out at the electrodes open circuit potential and utilised a 50 mV sinusoidal potential ranging from 10,000 Hz to 1 Hz.

### In vivo biocompatibility experiments

Ten female Sprague-Dawley rats, between 210 and 258 grams at time of surgery, were obtained from the Vernon Jansen Unit, University of Auckland in compliance with the University of Auckland Animal Ethics Committee Guidelines (AEC002056) and the New Zealand Animal Welfare Act 1999. Animals were housed together in a dedicated colony and behavioural room on a 12:12 light / dark cycle and always had ad libitum access to rat chow and water. They were group housed prior to surgery, after which they were individually housed.

#### Surgical procedure

Animals were anesthetised with isoflurane, temperature monitored and placed on a heating pad and given subcutaneous antibiotics (Baytril 100 mg/mL, 0.1 mL/100g) and analgesics (Carprofen 50 mg/mL, 0.01 mL/100g; Bupivicaine 2.5 mg/mL 0.16mL/100g; Buprenorphine 0.3 mg/mL, 0.17 mL/100g). A laminectomy was performed to expose the spinal cord at the spinal processes T10, T11, and T12. Small incisions in the dura were made near either end of the exposed spinal cord. A guidance catheter, containing a suture anchored to the tail of the implant, was inserted below dura at the T10 end and carefully propelled along the surface of the spinal cord, exiting at the T12 end. The suture was then used to guide the implant along the subdural space and used to position the implant so that the bioelectronic head was in contact with the spinal cord. The bifurcated design of the bioelectronic used two guidance catheters (one attached to either arm) which were used to guide and position the arms either side of the posterior spinal cord vein. Once the implant was positioned, the guidance suture was removed. A thin piece of absorbable gelatin sponge (Pfizer Gelfoam, Amtech Medical Limited, Wanganui) was used to form a haemostatic seal in the space above the exposed spinal cord. At the tail end, the implant was connected to a small PCB which routed the electrode channels to a 36-channel Omnetics female nano-strip plug. In some animals, the implant was briefly connected via a recording cable to MEA2100 system to make test recordings. The protruding section of implant near the printed circuit board was then severed. The deeper muscle tissue was sutured above the Gelfoam using an absorbable suture (ChomicGut 4.0) and the skin was sutured closed using nylon suture (Ethicon 4.0, Johnson and Johnson) so that no part of the implant was subcutaneous or external to the animal. This directly tested any effects of the presence of the implant along the spinal cord without any effects of additional forces associated with an external connector on the implant.

#### Post-operative Care

Post-operatively, animals were given subcutaneous antibiotics (Baytril 100 mg/mL, 0.1 mL/100g), analgesics (Carprofen 50 mg/mL, 0.01 mL/100g; Buprenorphine 0.3 mg/mL, 0.17 mL/100g), and supplementary fluid (Saline 3 mL) twice daily for 3 days. Bladder function was assessed, however, it was not necessary to manually void the bladder on any of the rats.

#### Assessment of locomotor function

The BBB locomotor rating scale was used to measure right and left hind limb motor function. Animals were placed in a circular open field (100 cm diameter white wooden floor with 20 cm perspex walls) in the recovery room. Following the surgery, rats were tested on post-operative day’s 1, 3, and 7 for 5 - 8 mins each time depending on the amount of movement exhibited. This length of time has been determined as providing sufficient time to observe and record behavioural recovery of individual rats with minimal risk of missing key findings. The two animals retained for 5 weeks were assessed with the BBB-scale on days 1, 3, 7, 14, 21, 28, and 35 post-surgery. BBB-scores were determined by a single experimenter (BH) blinded to the condition of the rats during scoring. The functionality of each hind-limb was scored from 0 (total paralysis) to 21 (normal movement) as described previously (*22*), and the scores for the right and left hindlimbs averaged.

#### Histological sectioning

At the end of the 1 or 5-week behavioural assessment period, animals were euthanised with sodium pentobarbitol (100 mg/kg, i.p.) and perfused intracardially with 400 mL of 0.9% Saline followed by 400 mL 4% paraformaldehyde in 0.1 M phosphate buffer (pH 7.4). Approximately 15 mm of spinal cord with the 10mm extent of spinal cord that contacted the bioelectronic head of the implant in the centre was dissected, removed and post fixed in 4% paraformaldehyde for four hours then transferred to 20% sucrose and stored at 4°C. After 3-4 days the cord was transferred to 30% sucrose containing 0.1% sodium azide. Spinal cords embedded in OCT compound were sectioned coronally (from caudal to rostral) at 10 μm using a Cryostat and mounted onto positively charged glass slides.

#### Haematoxylin and eosin staining

The majority of slides were stained with a haematoxylin and eosin (H&E) stain to aid with visualisation of the tissue for the assessment of shape and surface area of the spinal cord. Briefly, sections were hydrated and then immersed in Hematoxylin solution, differentiated in acid alcohol, colour-shifted towards blue using Lithium Carbonate, and then cross-stained with Eosin Y solution. Between these steps, slides were placed under running tap water for several minutes. Slides were then rehydrated using graded ethanols, deparaffinised in xylene and mounted and coverslipped using DPX.

#### Immunohistochemical staining

The remaining slides were stained with two standard cellular markers for foreign-body response of astrocytes (GFAP, red) and microglia (Iba1, green). Immunohistochemistry was carried out using the following antibodies; mouse anti-glial fibrillary acidic protein (GFAP)–Cy3 (Sigma, 1:2000) to show astrocyte structure, and goat anti-Iba1 (Abcam, ab5076, 1:250) to label microglia. Antibodies were diluted with 4% normal donkey serum (NDS) in PBS–T and incubated overnight at 4 °C. Control sections on each slide were GFAP only, Iba1 only, and NDS only (no primary antibody). The next day, slides were washed in PBS and incubated with a 1:250 dilution of donkey anti-goat Alexa-488 (Iba1) in 4% NDS in PBS–T for 4 h at room temperature. The nuclei of cells were counterstained using Hoechst 33342 (Thermo-Fisher Scientific #62249). Slides were washed with PBS and coverslip mounted using ProLong Gold Antifade Reagent (Invitrogen). Staining was done in batches, which each included a mix of slides from different experimental animals and groups.

#### Imaging and Analysis

Tiled imaging of whole H&E-stained coronal sections was performed using a 10x objective on an EVOS M7000 microscope. Immunohistochemically-stained whole sections were imaged using a 4x objective on an EVOS M5000 microscope to ensure no distortion of fluorescent intensity due to tiling. Fluorescent images were taken using the appropriate filter sets (RFP, GFP, DAPI) and all microscope settings were kept consistent. Imaged sections were aligned using the midline as an anatomical landmark and the filenames blinded by a colleague who was not part of the study. A maximum of three sections per spinal cord block were imaged per slide. Each coronal section was cross-referenced with a Rat Spinal Cord Atlas (*23*) to determine vertebral location along the spinal cord, and Harrison et al. (2013) was used to determine the corresponding spinal cord segment. The perimeter of each H&E-stained section was traced in ImageJ and the surface area (cm2) and shape (‘roundness’) measured. Roundness used the formula: 4 * area / (π * major_axis ^ 2). A value of 1.0 indicated a perfect circle, whereas decreasing values indicated an increasingly elongated shape with a value of 0 indicating a flat horizontal line (see **Figure 3D**, bottom left). For immunohistochemically-stained sections, analysis was restricted to a region of interest of tissue directly below the implant position (2.4 mm centred across the dorsal surface and 200 microns deep; see **Figure 4A**). The mean pixel intensity of fluorescence in this region was determined in ImageJ for the GFAP and Iba1 images from each section.

### Electrophysiological recording experiments

Four female Sprague-Dawley rats, between 235 and 241 grams at time of surgery, were obtained and housed in the same room and conditions as described previously.

#### Bioelectronic backpack assemblies

The polyimide implants were carefully soldered to pads on a small custom printed circuit board (10 x 12 mm) which routed the electrodes, two grounds, and reference to a 32-channel Omnetics female nanostrip connector soldered into drill hole vias on the other side and end of the board. Fine silk suture (Ethicon, 7.0) was inserted into a 1.5 cm piece of 28G polyurethane rat intrathecal catheter (Alzet), looped through an ear-hole at the tip of the implant arm, inserted back through the catheter, and then snipped and superglued at the tip of the catheter. The backpack was 3D printed in grey resin using a Creality LD-002R Resin 3D printer at 50 μm resolution. A ~ 2 cm diameter section of surgical mesh was attached to the ventral surface of the backpack with non-absorbable nylon suture (Ethicon 4.0) through 4 small slots in the midline body of device and through the hole at the apex of each of the five legs (see **Figure 6A**). The polyimide implant was carefully dropped through a small internal slot in the backpack and the PCB board and Omnetics connector positioned within the square upper section of the backpack. The components were then secured with a lid inserted into grooves on the dorsal surface and filled with epoxy resin glue. Electrochemical impedance spectroscopy (as described above) was used to test whether each electrode channel was valid, in which case it had an impedance of ~ 5.4 – 5.6 MΩ (at 1,000 Hz).

#### Surgical procedure

In these animals, bifurcated bioelectronic implants were inserted as described above. Once the gelfoam had been inserted and the deeper muscle sutured closed, the backpack and surgical mesh were positioned in a pocket beneath the skin just above the cranial end of the incision and in front of the laminectomy site. The body of the bioelectronic implant protruded from the deeper muscle below the surgical mesh at the rostral end of the assembly and was carefully folded back on itself below the assembly to provide slack. The five legs protruding from the ventral surface of the backpack were then sutured to the deeper muscle through the surgical mesh using non-absorbable nylon suture (Ethicon 4.0). Over several weeks, the mesh provided a scaffold for tissue regrowth resulting in a strong anchor ensuring minimal risk of the implant or backpack becoming detached. The skin was sutured closed tightly around the base of the backpack using silk suture (Ethicon 4.0, Johnson and Johnson), and the hind-limb claws were carefully trimmed under the microscope to reduce damage caused by scratching.

#### Electrical recording sessions

Rats were acquired 3-weeks before the surgery date, over which they were habituated to handling by the experimenters for at least 5 days per week. After several days of handling, the rats were habituated to sitting and ambulating upon a raised platform (130 cm height, ~ 32 cm diameter) over several weeks. During post-surgery recording sessions, rats were placed on the platform with the recording cable hanging overhead. The backpack was then gently grasped by an experimenter to aid in connection (and removal) of the recording cable plug. Recordings were then made for a duration of 3 – 5 mins depending on the composure of the animal.

#### Recording equipment and signal processing

Electrical recordings were acquired with a Multi-Channel Systems ME2100 (Harvard Bioscience) electrophysiology rig. The raw analog signals were amplified ten-times and acquired with a sampling frequency of 20 kHz per channel, which was converted to a digital signal at a resolution of 24 bits. Raw data was converted to HDF5 format and then further processed in MATLAB (2018b, Mathworks) using in-house scripts. Line noise from the power supply was removed using a 50 Hz notch filter. A median channel was then calculated and subtracted from each individual channel. The effect of the median channel subtraction can be seen in **Supplemental Figure 1**. Specifically, the median channel contains changes in voltage that are common to each electrode, and therefore likely artifactual. By subtracting the median, the processed data reflects changes in voltage that are unique to an individual, or a small number, of electrodes. The median subtraction was done separately for the electrodes of each arm of the implant. Individual voltage waveforms were manually segmented from the processed data. The start of each waveform was defined as the time of the first change in voltage, relative to the baseline, that was 10 % of the maximum initial voltage deviation. The end of each waveform was defined by the time the voltage returned to within 10 % of the baseline voltage, after any period of hyperpolarization. The baseline for single voltage spikes (**Figure 8, B1 – B2**) was calculated as the mean voltage between 0 – 200 ms prior to the start of the waveform. A time varying baseline was calculated for oscillatory voltage spikes (**Figure 8, B3**) according to the method defined in Jia et al. (*40*)

## Acknowledgements

This research was supported through research grants from the Neurological Foundation of New Zealand (1941 PG), the CatWalk Spinal Cord Injury Trust and the Health Research Council of New Zealand (HRC/Catwalk Partnership 19/895) and DS HRC Hercus Fellowship (19/007). Author’s BH and ZA contributed equally to this work.

## Author contributions

Conceptualization, Z.A., M.A., S.J.O. and D.S.; Methodology, B.H., Z.A., M.V., C.B., B.R., M.A., S.J.O, D.S.; Software, B.H., Z.A., B.R.; Formal Analysis, B.H., Z.A., B.R.; Investigation, B.H., Z.A., M.V., C.B., B.R.; Resources, B.H., B.R., M.A., S.J.O, D.S.; Writing – original draft, B.H., Z.A.; Writing Review & Editing, B.H., Z.A., M.V., C.B., B.R., M.A., S.J.O, D.S.; Visualization, B.H., B.R.; Supervision M.A., S.J.O, D.S.; Funding Acquisition B.H., Z.A., B.R., M.A., S.J.O, D.S.

## Supplemental Text and Figures

**Supplemental Figure 1:**
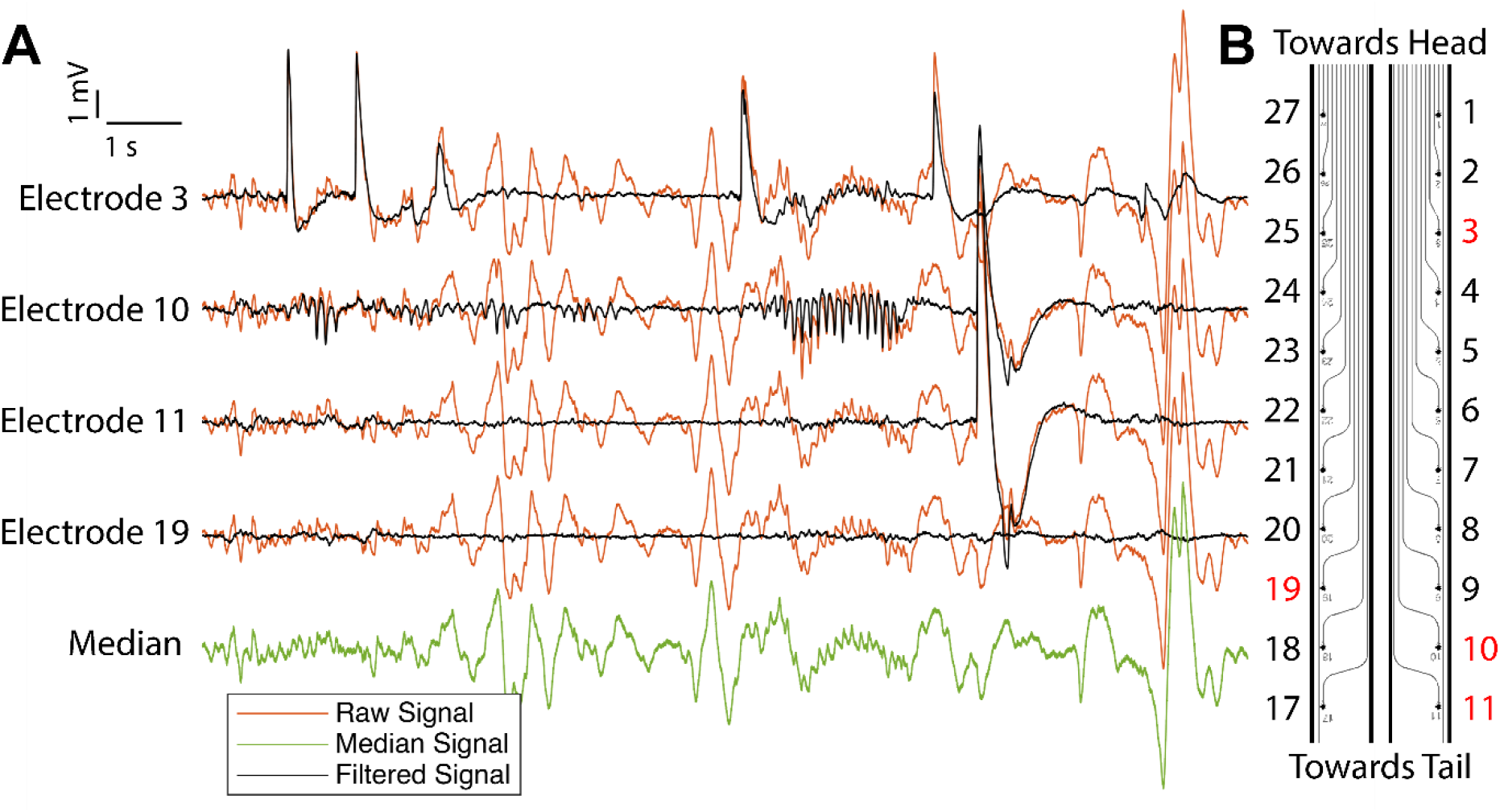
The effect of median channel subtraction on the electrical recordings. Data corresponds to electrodes 3, 10, 11 and 19, shown in Figure 7, between 214 – 220 seconds. (**A**) Raw, median and filtered signals for four individual electrodes, highlighted in (**B**) in red. The median channel contains changed in voltage that are common to all channels. Subtracting the median channel from each individual channel reveals electrical activity that is unique to a single or a small number of individual electrodes.

## References

1. C. Alvarado-Rojas et al., Single-unit activities during epileptic discharges in the human hippocampal formation. Frontiers in computational neuroscience 7, 140 (2013).

2. N. Cho, J. W. Squair, J. Bloch, G. Courtine, Neurorestorative interventions involving bioelectronic implants after spinal cord injury. Bioelectronic Medicine 5, 10 (2019).

3. G. A. Goetz, D. V. Palanker, Electronic approaches to restoration of sight. Rep Prog Phys 79, 096701–096701 (2016).

4. P. S. Olofsson, K. J. Tracey, Bioelectronic medicine: technology targeting molecular mechanisms for therapy. J Intern Med 282, 3–4 (2017).

5. C. Hassler, R. P. von Metzen, P. Ruther, T. Stieglitz, Characterization of parylene C as an encapsulation material for implanted neural prostheses. Journal of Biomedical Materials Research Part B: Applied Biomaterials 93B, 266–274 (2010).

6. T. Stieglitz, J. U. Meyer, Implantable microsystems. Polyimide-based neuroprostheses for interfacing nerves. Med Device Technol 10, 28–30 (1999).

7. E. M. Thaning, M. L. Asplund, T. A. Nyberg, O. W. Inganäs, H. von Holst, Stability of poly(3,4-ethylene dioxythiophene) materials intended for implants. Journal of biomedical materials research. Part B, Applied biomaterials 93, 407–415 (2010).

8. M. Capogrosso et al., A brain–spine interface alleviating gait deficits after spinal cord injury in primates. Nature 539, 284–288 (2016).

9. I. R. Minev et al., Electronic dura mater for long-term multimodal neural interfaces. Science 347, 159–163 (2015).

10. F. B. Wagner et al., Targeted neurotechnology restores walking in humans with spinal cord injury. Nature 563, 65-+ (2018).

11. C. Boehler et al., Actively controlled release of Dexamethasone from neural microelectrodes in a chronic in vivo study. Biomaterials 129, 176–187 (2017).

12. N. Wenger et al., Spatiotemporal neuromodulation therapies engaging muscle synergies improve motor control after spinal cord injury. Nat Med 22, 138–145 (2016).

13. M. Vomero et al., Conformable polyimide-based μECoGs: Bringing the electrodes closer to the signal source. Biomaterials 255, 120178 (2020).

14. K. K. Gokoffski, X. Jia, D. Shvarts, G. Xia, M. Zhao, Physiologic Electrical Fields Direct Retinal Ganglion Cell Axon Growth In Vitro. Invest Ophthalmol Vis Sci 60, 3659–3668 (2019).

15. S. Han et al., Electrical stimulation accelerates neurite regeneration in axotomized dorsal root ganglion neurons by increasing MMP-2 expression. Biochemical and biophysical research communications 508, 348–353 (2019).

16. M. D. Tang-Schomer, 3D axon growth by exogenous electrical stimulus and soluble factors. Brain Res 1678, 288–296 (2018).

17. J. M. Griffin et al., Astrocyte-selective AAV-ADAMTS4 gene therapy combined with hindlimb rehabilitation promotes functional recovery after spinal cord injury. Experimental Neurology 327, 113232 (2020).

18. C. Boehler, S. Carli, L. Fadiga, T. Stieglitz, M. Asplund, Tutorial: guidelines for standardized performance tests for electrodes intended for neural interfaces and bioelectronics. Nature Protocols 15, 3557–3578 (2020).

19. C. Boehler, T. Stieglitz, M. Asplund, Nanostructured platinum grass enables superior impedance reduction for neural microelectrodes. Biomaterials 67, 346–353 (2015).

20. Z. Aqrawe et al., The influence of macropores on PEDOT/PSS microelectrode coatings for neuronal recording and stimulation. Sensors and Actuators B: Chemical 281, 549–560 (2019).

21. Z. Aqrawe, J. Montgomery, J. Travas-Sejdic, D. Svirskis, Conducting polymers for neuronal microelectrode array recording and stimulation. Sensors and Actuators B: Chemical 257, 753–765 (2018).

22. D. M. Basso, M. S. Beattie, J. C. Bresnahan, A Sensitive and Reliable Locomotor Rating Scale for Open Field Testing in Rats. Journal of Neurotrauma 12, 1–21 (1995).

23. P. G. Watson C., Kayalioglu G., Heise C., in The Spinal Cord 1st Edition, P. G. Watson C., Kayalioglu G., Ed. (Academic Press, 2009), chap. 15, pp. 238–306.

24. A. Ersen, S. Elkabes, D. S. Freedman, M. Sahin, Chronic tissue response to untethered microelectrode implants in the rat brain and spinal cord. J Neural Eng 12, 016019 (2015).

25. P. Moshayedi et al., The relationship between glial cell mechanosensitivity and foreign body reactions in the central nervous system. Biomaterials 35, 3919–3925 (2014).

26. V. S. Polikov, P. A. Tresco, W. M. Reichert, Response of brain tissue to chronically implanted neural electrodes. J Neurosci Methods 148, 1–18 (2005).

27. Z. Aqrawe et al., A simultaneous optical and electrical in-vitro neuronal recording system to evaluate microelectrode performance. PloS one 15, e0237709 (2020).

28. G. K., in The Spinal Cord 1st Edition, P. G. Watson C., Kayalioglu G., Ed. (Academic Press, 2009), chap. 10, pp. 148–167.

29. M. E. Larkum, M. G. Rioult, H. R. Luscher, Propagation of action potentials in the dendrites of neurons from rat spinal cord slice cultures. Journal of Neurophysiology 75, 154–170 (1996).

30. G. Zhong et al., Electrophysiological Characterization of V2a Interneurons and Their Locomotor-Related Activity in the Neonatal Mouse Spinal Cord. The Journal of Neuroscience 30, 170 (2010).

31. P. M. Klinge et al., Immunohistochemical characterization of axonal sprouting and reactive tissue changes after long-term implantation of a polyimide sieve electrode to the transected adult rat sciatic nerve. Biomaterials 22, 2333–2343 (2001).

32. B. Rubehn, T. Stieglitz, In vitro evaluation of the long-term stability of polyimide as a material for neural implants. Biomaterials 31, 3449–3458 (2010).

33. Y. Sun et al., Assessment of the biocompatibility of photosensitive polyimide for implantable medical device use. Journal of biomedical materials research. Part A 90, 648–655 (2009).

34. G. Schiavone et al., Guidelines to Study and Develop Soft Electrode Systems for Neural Stimulation. Neuron 108, 238–258 (2020).

35. D. Mazensky, S. Flesarova, I. Sulla, Arterial Blood Supply to the Spinal Cord in Animal Models of Spinal Cord Injury. A Review. Anat Rec (Hoboken) 300, 2091–2106 (2017).

36. S. Wurth et al., Long-term usability and bio-integration of polyimide-based intra-neural stimulating electrodes. Biomaterials 122, 114–129 (2017).

37. B. Harland, M. Contreras, M. Souder, J.-M. Fellous, Dorsal CA1 hippocampal place cells form a multi-scale representation of megaspace. Current Biology, (2021).

38. B. Harland et al., Lesions of the Head Direction Cell System Increase Hippocampal Place Field Repetition. Curr Biol 27, 2706–2712 e2702 (2017).

39. W. D. Dietrich, Protection and Repair After Spinal Cord Injury: Accomplishments and Future Directions. Top Spinal Cord Inj Rehabil 21, 174–187 (2015).

40. H. Jia, N. L. Rochefort, X. Chen, A. Konnerth, In vivo two-photon imaging of sensory-evoked dendritic calcium signals in cortical neurons. Nature Protocols 6, 28–35 (2011).

